# Uroprotective effect of a protein isolated from seed of *Morinda citrifolia* (McLTP_1_) on hemorrhagic cystitis induced by ifosfamide in mice

**DOI:** 10.1101/2023.01.30.526355

**Authors:** Gisele de Fátima Pinheiro Rangel, Aurilene Gomes Cajado, Anamaria Falcão Pereira, Liviane Maria Alves Rabelo, Andrea Santos Costa, Hermógenes David de Oliveira, Deysi Viviana Tenazoa Wong, Renata Ferreira de Carvalho Leitão, Nylane Maria Nunes de Alencar

## Abstract

Hemorrhagic cystitis is a side effect of chemotherapy induced by an antineoplastic agent from the oxazaphosphorine group (ifosfamide and cyclophosphamide), resulting from the formation of the urotoxic metabolite acrolein. Morinda citrifolia Linn., popularly known as noni, is a species of Rubiaceae, where it is used from the root to the fruit for therapeutic purposes. From the seeds, a thermostable protein called McLTP_1_ (9.4 kDa) was extracted, among its therapeutic effects, it showed anti-inflammatory, gastroprotective, antibacterial and antinociceptive activity. Thus, the objective of this study is to evaluate the protective effect and the possible mechanism of action of a protein isolated from the seed of Morinda citrifolia (McLTP_1_) in hemorrhagic cystitis induced by ifosfamide in mice. Hemorrhagic cystitis was induced by intraperitoneal (i.p) administration of ifosfamide (IFO) in a single dose of 400mg/kg, according to a standardized protocol, in male balb/c mice. The experimental group treated with the uroprotective drug, mesna (80 mg/kg; i.p), received a pretreatment 30 minutes before, 4 and 8 hours after IFO. Treatment with McLTP_1_ was divided into two protocols, the first to define the best dose through a dose-response curve, where a pre-treatment was performed three days before cystitis induction, with McLTP_1_ administered at doses of 10, 20 or 40mg/kg (i.p), and two treatments 2 and 4 hours after IFO administration, evaluating its effect on bladder wet weight, edema and hemorrhage scores, and neutrophilic infiltrate. In the second protocol, only the best dose was used for the analysis of its effect on the hemorrhagic cystitis model. After 12 hours of hemorrhagic cystitis induction, the animals were euthanized by a high anesthetic dose. Subsequently, the bladders were removed, weighed and kept in 10% buffered formalin for histological, immunohistochemical (COX-2 and TNF-α), immunofluorescence (NF-kB and F4-80) analyses, or stored at -80°C for of MPO, vascular permeability, hemoblobin, cytokines (TNF-α, IL-1β, IL-6, IL-10, IL-4, IL-33), enzymes (iNOS and COX-2) and markers of oxidative stress (MDA, NO, GSH, SOD and CAT). The adopted experimental procedures were approved by the Animal Research Ethics Committee through protocol number 23170920-0. Treatment with McLTP1 reduced bladder wet weight at the three respective doses mentioned above, however, it was observed the reduction of toxicity parameters (macroscopic edema and hemorrhage scores) only at the lowest dose (10 mg/kg), as well as MPO activity at doses of 10 and 20 mg/kg (p<0.05). results, the lowest dose was chosen for subsequent results. McLTP_1_ (10 mg/kg) was able to promote permeability reduction and vascular and hemoglobin in the bladder through quantification by the evans blue method and cyanmethemoglobin, respectively (p<0.05). In addition, it had a protective effect by attenuating inflammatory scores and preserving the structure of the urothelium. The anti-inflammatory activity was demonstrated through the significant decrease of the cytokines TNF-α, IL-1β, IL-6 and increase of IL-10; reduced expression of COX-2, NF-kB and F4/80, and gene expression of IL-33, IL-4 and iNOS (p<0.05). McLTP_1_ also showed antioxidant activity, being able to reduce MDA and NO and increase levels of GSH, SOD and CAT (p<0.05). From the presented data, we can infer that McLTP_1_ is a potential uroprotector in the prevention of ifosfamide-induced hemorrhagic cystitis in mice by reducing inflammatory parameters and antioxidant activity.

## Introduction

Ifosfamide, a alkylating chemotherapy agents, is used in a wide diversity of malignancies including soft tissue sarcomas, leukemias, and bladder, testis, lung cancer [1]. Ifosfamide is primarily metabolized in the liver into acrolein, which is excreted in the urine and causing a deleterious effect on the urothelium [2]. Hemorrhagic cystitis is an inflammatory condition of the bladder that results in mucosal bleeding associated with hematuria manifesting as a primary clinical sign. The pathophysiology of this adverse effect involves oxidative stress, inflammation, and nitric oxide release, resulting in bladder hemorrhage, edema, urothelial denudation, and infiltration of inflammatory cells [3]. Management of ifosfamide hemorrhagic cystitis can lead to the need for clot extraction by cystoscopy, continuous bladder irrigation, and even intravesical drug administration. In severe cases, it can lead to the need for blood transfusion and even cystectomy [4].

Oxazaphosphorine-induced bladder inflammation is mediated by pro inflammatory cytokines, including such as interleukin-1β (IL-1β), tumor necrosis factor-α (TNF- α),, interleukin-6 (IL-6), cyclooxygenase-2 (COX-2), nitric oxide (NO) and NF-kB, which cause edema and hemorrhage associated with epithelial damage in the tissue [5–9]. These mediators partially contribute to bladder damage via the recruitment of neutrophils, since inactivation of these inflammatory markers reduces neutrophil accumulation in the bladder and consequently the severity of hemorragic cystitis [10]. Acrolein is also accountable for the activation of intracellular reactive oxygen species and the formation of nitric oxide, leading to the increase of peroxynitrite, a potent oxidant which culminates in lipid peroxidation. Overproduction of the reactive oxygen species (ROS) and RNS during inflammation leads to cellular damage, including DNA injury, cellular proteins denaturation, and membranes lipid peroxidation [11,12].

One of the chemoprotective agents is mesna (2-mercaptoethanesulfonate) which directly binds, neutralizes acrolein and thus blocks its entry into the bladder cells thus preventing inflammation and oxidative stress [13,14]. However, mesna it is not effective to treat established hemorrhagic cystitis [15,16]. This occurs because of the other substances that do not bind to mesna and that have the capacity to cause hemorragic cystitis [17]. Therefore, diverse natural and synthetic substances has been investigated, such as anti-inflammatory drugs and antioxidants seems to be one of the interesting strategies in cases of inflammatory bladder conditions refractory to the conventional treatment [18–22].

*Morinda citrifolia* L. (Rubiaceae), popularly known as noni, is a plant native to tropical Asia, which has been used in popular medicine to the treatment of several diseases, such as hypertension, diabetes, rheumatoid arthritis, inflammation, and pain management [23]. In a pioneering way, Campos et al. (2016) isolated a thermostable to the lipid transfer protein class, named McLTP_1_, isolated from *Morinda citrifolia* L. seeds (UniProtKB accession number: C0HJH5), exhibited anti-inflammatory and antinociceptive activities when orally administered to mice, besides gastroprotective, antibacterial and nephroprotective investigated in different experimental models. Hence, the aim of this study was to evaluate the uroprotective effect of a protein isolated from the seed of Morinda citrifolia (McLTP1) on ifosfamide-induced hemorrhagic cystitis in mice, investigating the mechanisms of action involved.

## Materials and methods

### Purification of McLTP1

McLTP_1_ was purified from defatted noni (Morinda citrifolia L.) seeds flour from as previously described by Campos et al. (2016) [24]. After isolation, the purity of McLTP1 was measured by non-reducing 15% sodium dodecyl sulfate-polyacrylamide gel electrophoresis (SDS-PAGE) [25]. McLTP1 samples used in biological assays were prepared based on soluble total protein concentrations (mg/ml) valued by the method of Bradford (1976) [26].

### Animals

Balb/c male mice (Mus musculus) (8 weeks old) from the Biotherium of the Nucleus for Research and Development of Medicines of the UFC were used to execute the experimental protocols (n = 6-8/group). The animals were maintained on a 12-hour light/dark cycle, temperature (23 ± 2 °C) and humidity (55 ± 10%) controlled, with ad libitum supply of food and water. The adopted experimental procedures were approved by the Animal Research Ethics Committee of the Research and of the Federal University of Ceará through protocol number 23170920-0.

### Ifosfamide°induced hemorragic cystitis and dose treatment schedule

Mice were randomly divided into six groups: (1) control group vehicle, (2) Ifosfamide (Eurofarma; Brazil; 400 mg/kg)-treated group, (3) IFO + Mesna (Mitexan®; 80 mg/kg) treated group, (4) IFO + McLTP_1_ (10 mg/kg) treated group, (5) IFO + McLTP_1_ (20 mg/kg) treated group, and (6) IFO + McLTP_1_ (40 mg/kg) treated group. In the control group, mice were injected with equivalent volumes of saline. In group 2, mice were pretreated with saline once a day for 3 consecutive days, followed by IFO (400 mg/kg) administration 30 min after the last dose of saline on the third day. In group 3, mice were treated with Mesna 30 min before and 4 and 8 h after cystitis induction with a single dose of IFO (400 mg/kg). In groups 4, 5 and 6; mice were pretreated with McLTP_1_ once a day for 3 consecutive days, with the respective doses mentioned above, followed by IFO (400 mg/kg) administration 30 min after the dose of McLTP_1_ on the third day, and 4 and 8 h after cystitis induction with IFO. All drugs were injected intraperitoneally. Drug doses and protocol were selected based on previous studies [18,20,27,28]. 12 h after IFO administration, the animals were euthanized by high anesthetic dose with xylazine hydrochloride (30 mg/kg, i.p.) and ketamine hydrochloride (300 mg/kg, i.p.). Then, the bladders were removed, weighed and placed in 10% buffered formalin or stored at -80°C for subsequent analysis.

### Bladder wet weight evaluation

Bladders were dissected, and the remaining urinary content was removed. Bladder wet weight (BWW) was geted, and the results are expressed in milligrams per 20 g of body weight [mean ± standard error of mean (SEM)], a parameter that represents the bladder edema.

### Macroscopic evaluation

The bladders were visually analyzed for edema and hemorrhage according to the Criteria of [29]: normal (0); intermediary (1+), between normal and moderate; moderate (2+), fluid confined to the internal mucosa; and intense (3+), fluid seen externally and internally in the bladder walls. Hemorrhage was scored as follows: normal (0), absence of hemorrhage and vessel dilation; intermediary (1+), telangiectasia or dilation of bladder vessels; moderate (2+), mucosal hemorrhage; severe (3+), intravesical clots [29].

### Myeloperoxidase (MPO) assay

Myeloperoxidase, an enzyme present in the azurophilic granules of neutrophils, is used as a quantitative marker of the accumulation of this cell in the inflammatory site. The bladder was homogenized using a grinder (Pollytron®) in 200μL of ice-cold buffer (0.1 M NaCl, 0.02 M NaPO_4_, 0.015 M NaEDTA; pH 4.7) and centrifuged at 3,000 rpm for 15 min at 4°C. The pellet was resuspended to hypotonic lysis (0.2% NaCl solution) followed by a further centrifugation step. The pellet was resuspended in 300 μL of 0.05M NaPO4 buffer (pH 5.4) with hexadecyltrimethylammonium bromide (HTAB, Sigma), then the material was macerated. The homogenate was centrifuged at 10,000 rpm for 15 min, 4°C.

Subsequently, 50 μL of the supernatant were applied to a 96-well plate for the assay. In each well, 25 μL of TMB (3, 3’, 3, 3-tetramethylbenzidine; 1.6 mM) and 100 μL of H_2_O_2_ (0.5 mM) were added and the plate was incubated in an oven for 5 min at 37 °C. Then, the reaction was stopped with 4M sulfuric acid. Quantification of neutrophils was performed using a standard neutrophil curve (with 1×10^5^ neutrophils/50 μL in the first well). The absorbances were read using a spectrophotometer at a wavelength of 450 nm, and the results were expressed as the number of neutrophils/mg of tissue [30].

### Evan’s Blue Extravasation Assay

The increase in vesical vascular permeability was evaluated through the extravasation of Evans Blue dye. The dye was injected intravenously via the retro orbital plexus at a dose of 25 mg/kg 30 minutes before the animals were euthanized. The bladders were removed, the urinary contents were discarded and they were incubated in formamide solution (1 ml/bladder) at 56°C overnight. The extracted dye was determined by measuring the change in absorbance at 630 nm in an ELISA reader. Vascular permeability was expressed in μg of Evans Blue/mg of tissue (mean ± SEM) [31].

### Hemoglobin Assay

The presence of hemorrhage in the bladder was determined by a hemoglobin assay using the cyanomethemoglobin method described by Harold and Drabkin (1935). The bladders were homogenized in Drabkin’s reagent (100 mg of bladder tissue per mL of reagent) and after 1 h of incubation, the bladders were centrifuged at 10,000 G for 10 min. Supernatants were extracted and centrifuged again at 10,000 G for 10 min. The absorbance of the supernatant was quantified using an ELISA reader at a wavelength of 540 nm and the hemoglobin concentration was calculated through the previously constructed analytical curve using a hemoglobin standard (Labtest, Brazil) and the results were expressed in μg of hemoglobin/mg of tissue [32].

### Microscopic Evaluation

The bladders were removed and fixed in 10% buffered formalin for 24 hours, then dehydrated in 70% alcohol and embedded in paraffin. Paraffin blocks were cut into 5µm and stained with hematoxylin-eosin (HE). Histopathological analysis was performed by a blinded person who was unaware of the treatments and group divisions, and scored by Modified Gray criteria [33]: Severe alterations (3+) Characterized by multiple mucosal ulcerations, mucosal erosion, intense edema, intense inflammatory infiltrate, intense fibrin deposits and several hemorrhagic foci with possible transmural hemorrhage. Moderate changes (2+) Characterized by multiple mucosal ulcerations, moderate edema, inflammatory infiltrate, fibrin deposits and hemorrhagic foci. Slight changes (1+) Characterized by the number of epithelial cells decreased by desquamation, “erasure” of the usual folds of the mucosa due to submucosal edema, slight bleeding and few ulcers. Normal histology (0) Characterized by normal urothelium, as well as absence of ulcer and inflammatory infiltrate.

### Evaluation of hemorrhage by Masson’s trichrome

Bladder hemorrhage was evaluated in the different experimental groups from tissue slides stained with Masson’s special stain, a dye commonly used to identify collagen fibers, which are highlighted in blue in this stain. The slides were used in the present study for the evaluation of blood vessels, considering that red blood cells and endothelial cells are strongly marked in red, which facilitates the identification of blood vessels and collagen [34].

### Measuring the thickness of the urothelium

The measurement of the thickness of the urothelium was performed using the software ImageJ version 1.36b, measuring five serial images of the slide of each animal in each group using low magnification (×10), the photos were obtained in the Nikon Microscope with 10x objective and ocular lens Leitz Wetzlar micrometer 10x. The measured value was presented in millimeters [35].

### Immunohistochemistry for TNF-α and COX-2

Bladder samples were fixed in 10% buffered formalin for 24h and processed for paraffin embedding. After embedding in paraffin, the segments were cut in a microtome, obtaining thicknesses of 4µm that were inserted in silanized histological slides. In the next step, the slides containing the sections were subjected to deparaffinization and rehydrated through xylene and graded alcohols. At the end of this step, antigen retrieval was performed with citrate buffer (DAKO, pH 6.0) and endogenous peroxidase was blocked with 3% hydrogen peroxide (DAKO) for 30 min. After this time, the slides were washed with PBS and incubated with primary antibodies COX-2 (Invitrogen) (1:200 dilution) or TNF-α (Abcan) (1:300 dilution) overnight. After this period, the sections were incubated with HRP polymer (DAKO) for 30 min. Then, antibodies was visualized using the chromogen 3,3′-diaminobenzidine (DAB). Finally, the slides were counterstained with Mayer’s hematoxylin and processed to insert the coverslip. Negative control sections were processed simultaneously as described above, but with the first antibody being replaced by PBS–BSA 5%. None of the negative controls showed immunoreactivity. Tissue images were captured using a digitized camera coupled to the microscope (Nikon Elipse E200), capturing 10 fields per histological section at 200x or 400x magnification to obtain the total area of the tissue and the immunostained area. Quantification was performed by quantifying pixels by greater color saturation related to immunostaining with DAB, using the ImageJ software.

### F4/80 and NF-KB detection by immunofluorescence

Bladder sections were deparaffinized for 1 h at 60 °C. The tissues were then rehydrated in alcohol solutions, and antigen retrieval was performed in 0.1 M sodium citrate buffer (pH 6.0) at 95 °C for 18 min. After nuclear membrane permeabilization with a 10-min exposure with 0.2% Triton X-100 (Sigma-Aldrich®, St. Louis, MO, USA) solution, nonspecific binding was achieved by using 0.3 M glycine (Sigma-Aldrich®, St. Louis, MO, USA) in 5% bovine serum albumin (BSA) (Sigma-Aldrich®, St. Louis, MO, USA) for 30 min. Tissues were incubated overnight at 2–8 °C with rabbit-produced anti NF-KB p65 primers, nuclear labeling antibody (Abcan) (dilution: 1:100) or anti-F4/80 (Invitrogen®, Life Technologies, Thermo Fisher Scientific, Waltham, MA, USA) (1:200 dilution). Subsequently, after three baths in PBS for 5 min, incubation was performed with the secondary antibody made in Alexa FluorTM 568 donkey anti-rabbit (Invitrogen®, Life Technologies, Thermo Fisher Scientific, Waltham, MA, USA), at a dilution of 1 : 200, for 1 hour and 30 minutes. In order to mark cell nuclei, tissue sections were incubated for 30 minutes with DAPI (Invitrogen®, Life Technologies, Thermo Fisher Scientific, Waltham, MA, USA) (4 µL in 200 mL of PBS). To mount the slides, Prolong Gold Antifade Mountant medium (Invitrogen®, Life Technologies, Thermo Fisher Scientific, Waltham, MA, USA) was used. For the acquisition of photomicrographs, a laser scanning confocal microscope (Zeiss LSM 710, Carl Zeiss, Jena, Germany) with significant “master gain” and “digital offset” was used for further analysis. Photomicrographs were transmitted through imaging software (Fiji Image J, National Institutes of Health, Washington, DC, USA). In order to quantify the fluorescent area in the photomicrographs, it was necessary to differentiate the fluorescent pixels by the greater color saturation related to fluorescence (red). Previously, for the purpose of defining the selected and unselected pixels, the lower and upper limits were highlighted by the color threshold. The results obtained were expressed in percentage, which were compared to positive fluorescence of F4/80 or NF-KB (100%).

### TNF-α, IL-1β, IL-6 and IL-10 enzyme-linked immunosorbent assay (ELISA)

The concentration of cytokines present in the bladders was quantified through enzyme immunoassay (ELISA) according to the previously described methods [36]. Tissues were homogenized in 10% PBS (pH 7.4) (mg of tissue/µL) and centrifuged at 5,000 rpm for 10 minutes to use the supernatant. High-binding surface 96-well plates were incubated overnight with murine capture antibodies to TNF-α, IL-1β, IL-6 and IL-10 (kits from R&D systems). After blocking nonspecific binding sites with BSA (1%), 100 µL of sample were incubated for 2h at 4°C, as well as the serial standard curve also added in this step. After each incubation step, the plates are washed with PBS/TWEEN-20 buffer, and then detection antibody was added for 2h. Subsequently, the plates were washed and 100 μL of the HRP-streptavidin complex diluted (1:40) was added. After 20 minutes, 100 μL of TMB substrate (3,3’,5,5’-tetramethylbenzidine) was added and the plates were incubated in the dark at room temperature for 20 min. The enzymatic reaction was interrupted with 50 μL of H_2_SO_4_ (2N) and then the absorbance reading was measured in a spectrophotometer using a wavelength of 450 nm. Cytokine concentrations were calculated from a standard curve. The results were expressed in picogram of cytokines/mg of tissue.

### IL-4, IL-33, COX-2 and iNOS expression by real-time PCR

Bladder samples were homogenized in 1 mL of Trizol, RNA extraction followed the Trizol method according to the manufacturer’s specifications. Then, the samples were quantified in a Take3™ Microvolume Plate, using an Epoch™ Microplate Spectrofotometer (BioTek), samples with a purity factor where the absorbance A260/280 ranged from 1.5 to 2.0 were considered adequate for the study. RNA transcription into complementary DNA was performed using the High Capacity cDNA Reverse Transcription Kit in Veriti 96 Thermocycler both from Applied Biosystems. Finally, qRT-PCR was performed using the PowerUp SyBr Green Master Mix on the QuantStudio™ 3 Real-Time PCR System (Applied Biosystems, Thermo Scientific). Amplification was performed using the rapid cycling mode following the manufacturer’s specifications.The primers were designed based on nucleic acid sequences obtained from the National Center for Biotechnology Information and the primer sequences are displayed in Supplementary Table S1. Samples were normalized using the β-actin gene. Relative gene expression was determined using the 2-ΔΔCt method (Livak & Schmittgen, 2001).

### Determination of the concentration of Malondialdehyde (MDA)

The determination of the levels of reactive acid substances with thiobarbituric acid were measured through the quantification of malondialdehyde, a product of the decomposition of hydroperoxides of polyunsaturated fatty acids, formed during the oxidative process, according to the method described [37]. The bladders were homogenized with 0.05 M 10% phosphate buffer (mg of tissue/μL), 250μL of the homogenate were incubated in a water bath at 37°C for 1 hour. Then, in order to interrupt the peroxidation process, 400 μL of 35% perchloric acid was added to the samples and then centrifuged at 14,000 rpm, 15 minutes, 4 °C. The supernatant was collected and 200 μL of 0.8% thiobarbituric acid were added and again submitted to a water bath for 30 minutes at 95°C. After cooling, the samples were plated and a standard curve with tetramethoxypropane (TMP) was used for method calibration. Absorbances were measured at a wavelength of 532 nm. Results were expressed in nmol MDA/mg tissue.

### Determination of nitrite (NO) levels

This method is an indirect way of quantifying nitric oxide, since the NO^3−^ ion (nitrate) is determined as NO^2−^ ion (nitrite) after reduction by the enzyme nitrate reductase [38]. Bladders were homogenized with 10% potassium chloride buffer (0.15 M KCl (µL/mg of tissue), centrifuged at 5,000 rpm for 30 minutes. Then, 80 μL of the supernatant was incubated with the reagent solution (40 μL of nitrate reductase enzyme, NADPH substrate, KH_2_PO_4_ in distilled water) overnight in an oven at 37 °C, so that all nitrate present in the supernatant was converted into nitrite. The standard curve was also plated from a sodium nitrate solution (NaNO_2_ 200 μM). Finally, 80 μL of Griess solution (1% suffanilamide in 1% H_3_PO_4_/0.1% NEED/distilled water/1:1:1:1) was added to each well. The absorbances were read at 540 nm and the results were expressed in μM of NO^2−^.

### Determination of reduced glutathione (GSH) concentration

The determination of reduced glutathione, a water-soluble antioxidant endogenous component of the pool of non-protein sulfhydryl groups (NPSH), was analyzed according to the method [39] which is based on the reaction of DTNB (5,5-dithiobis acid 2-nitrobenzoic acid) with sulfhydryl compounds. The bladders were homogenized at 10% with 0.02 M EDTA buffer. Next, 60 μL of 10% trichloroacetic acid (TCA) was added to 40 μL of sample and then centrifuged at 5000 rpm, 15 min, 4°C. The supernatant was plated, as well as the standard curve with reduced glutathione. Finally, 102 μL of the reading solution (Tris-EDTA, DTNB 0.01 M) was added and the absorbance was measured at a wavelength of 412 nm. The results were expressed in µg of NP-SH/mg of tissue.

### Determination of catalase enzymatic activity (CAT)

Catalase is an antioxidant capable of transforming hydrogen peroxide (H_2_O_2_) into water and oxygen, the test determines the speed of consumption of H_2_O_2_ by the enzyme present in the sample, as described [40]. Bladders were homogenized at 10% in 150 mM phosphate buffer and then centrifuged at 10,000 rpm, 5 minutes, 4°C. And then, 20 μL of the supernatant were added to 980 μL of the reaction medium (0.1% H_2_O_2_, 5 mM Tris-HCl-EDTA pH 8.0 and Milli-Q water). The absorbances were measured after 1 and 6 minutes of addition of the reaction medium at a wavelength of 230 nm. Results were expressed as catalase units/mg of protein.

### Determination of the enzymatic activity of superoxide dismutase (SOD)

Superoxide dismutase (SOD) is an antioxidant enzyme that catalyzes the dismutation of superoxide anion to hydrogen peroxide (H_2_O_2_), a less harmful radical, which is then degraded by catalase. SOD is able to inhibit the reduction of nitro-tetrazolium blue (NBT) formazam by superoxide anion (O2-), in this methodology the photochemically reduced riboflavin releases O2-. Bladders were homogenized at 10% in 50 mM phosphate buffer (pH 7.8) and then centrifuged at 12,000 rpm, 20 min, 4°C. Of the obtained supernatant, 50 μL were added to 1 mL of the reaction medium (50 mM phosphate buffer, 0.1 mM EDTA, 19.5 mM methionine), 300 μL of 5 μM riboflavin and 150 μL of 750 μM NBT, protected from light. The material was exposed to fluorescent light (15 W) for 15 minutes and then plated for absorbance reading at a wavelength of 560 nm. The results were expressed in enzyme units (U)/mg of protein [41].

### Statistical analysis

Statistical analysis was performed using GraphPadPrism® software, version 8.0. The multiple comparison test was performed using analysis of variance (ANOVA) followed by the Bonferroni test (parametric data) or application of the Kruskal-Wallis test followed by the Dunn’s post-test (non-parametric data). Results were expressed as mean ± mean standard error (S.E.M.) or median, maximum, and minimum value. Results were considered statistically significant at P<0.05.

## Results

### Effect of different doses of McLTP_1_ in bladder wet weight, macroscopic scores and vesical neutrophil infiltration on ifosfamide-induced hemorrhagic cystitis

Hemorrhagic cystitis was induced by administering ifosfamide at a dose of 400 mg/kg intraperitoneally. IFO significantly increased bladder wet weight (39.61 ± 1.05 mg/20g of the animal body weight), Fig. 1A) and neutrophil accumulation (3730 ± 115.8 neutrophils/mg of tissue, Fig. 1B) compared to the saline group (bladder wet weight: 17.98 ± 0.87 mg/20g; neutrophil infiltration: 2754 ± 62.64). In addition, McLTP_1_ (10, 20 and 40 mg/kg) significantly attenuated IFO-induced bladder edema (30.24 ± 1.7; 28.29 ± 1.96; 26.89 ± 2.62 mg/20 g body weight, respectively, Fig. 1A) and prevented neutrophil accumulation at the doses 10 and 20 mg/kg (2491 ± 154.9; 2862 ± 157.5 neutrophils/mg of tissue, Fig. 1B) compared to the IFO group. As well as the mesna group (bladder wet weight: 29.22 ± 1.48; neutrophil infiltration: 2166 ± 74.09). However, the dose of 40ml/kg did not present a significant reduction (2928 ± 276.6 neutrophils/mg of tissue)

**Fig. 1.**
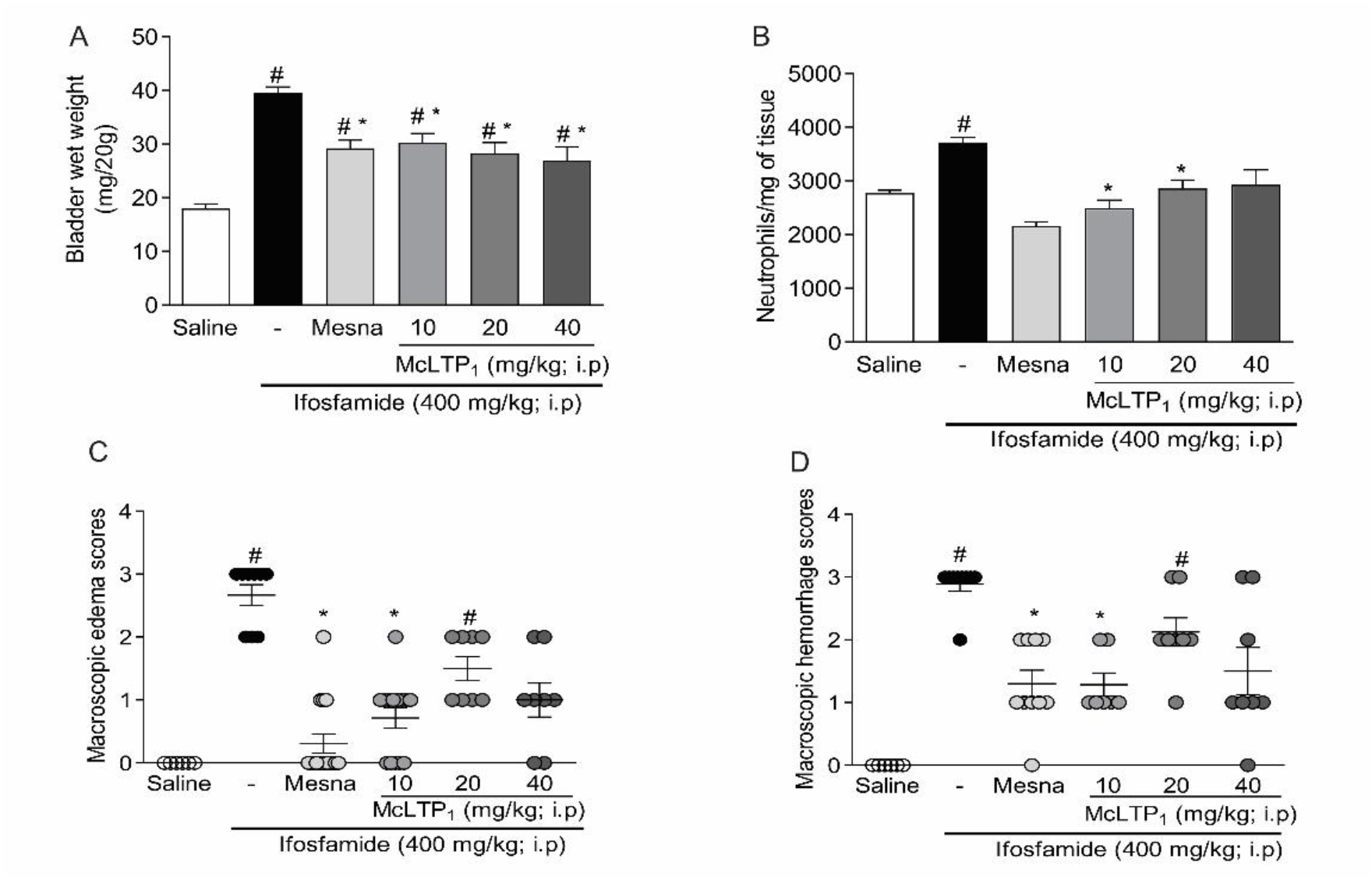
Effect of McLTP_1_ on bladder edema, macroscopy injury and neutrophil infiltrate during ifosfamide-induced hemorrhagic cystitis. The results are reported as means ± EPM (n = 6-8/group). #P < 0.05 vs. saline group (negative control); *P < 0.05 vs. IFO group. The results are reported as medians (minimum-maximum) (n = 6-8 per group) and analyzed using Kruskal-Wallis and Dunn’s test. #P < 0.05 vs. saline; *P < 0.05 vs. IFO.

IFO induced significant macroscopic (edema: 3[2-3] and hemorrhage: 3[2-3]) and compared to the saline group (edema: 0[0–0]; hemorrhage: 0[0–0]. Such damage was prevented in animals treated with McLTP1 (10 mg/kg) (edema: 1[0-2]; hemorrhage: 1[1–2]; compared to IFO. All of these parameters were also attenuated by mesna, which was used as a positive control [E: 0 (0-2); H: 1(0-2)]. However, at doses of 20 [E: 1.5 (1-2); H: 2 (1-3)] and 40mg/Kg [E: 1 (0-2); H: 1 (0-3)] there was no significant reduction (Fig 1C e 1D). The above results show that the lowest dose with the best therapeutic effect of McLTP1 is 10mg/kg, thus being eligible for carrying out the experiments below.

### Effect of McLTP1 on vascular permeability and hemoglobin dosage in ifosfamide-induced hemorrhagic cystitis

As shown in Fig. 2A, IFO significantly increased vascular permeability (0.32 ± 0.02 μg/mg of tissue) compared to the saline group (0.07 ± 0.004) and McLTP1 significantly reduced this parameter compared to the IFO group (0.20 ± 0.05). However, treatment with mesna did not significantly reduce this parameter when compared to the ifosfamide group (0.20 ± 0.05). In contrast to the saline group (9.79 ± 1.12 hemoglobin/mg of bladder), the bladder wall in animals treated with ifosfamide showed a significant increase in hemorrhage (22.40 ± 1.32), as indicated in Fig. 2B. McLTP1 treatment reduced the amount of hemoglobin (12.27 ± 0.42), similar to mesna treatment (13.08 ± 1.44).

**Fig. 2.**
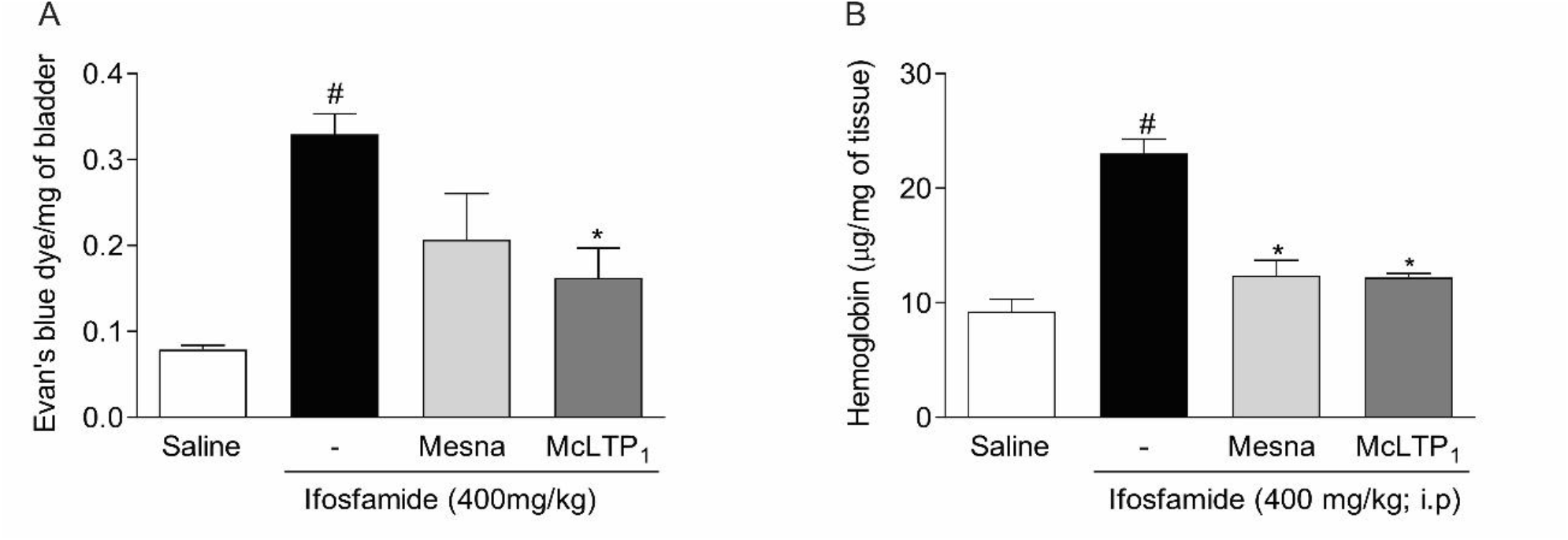
Effect of McLTP_1_ on the parameters of hemorrhage and tissue edema during ifosfamide-induced hemorrhagic cystitis. The results are reported as means ± EPM (n = 6-8/group). #P < 0.05 vs. saline group (negative control); *P < 0.05 vs. IFO group.

**Fig. 2.**
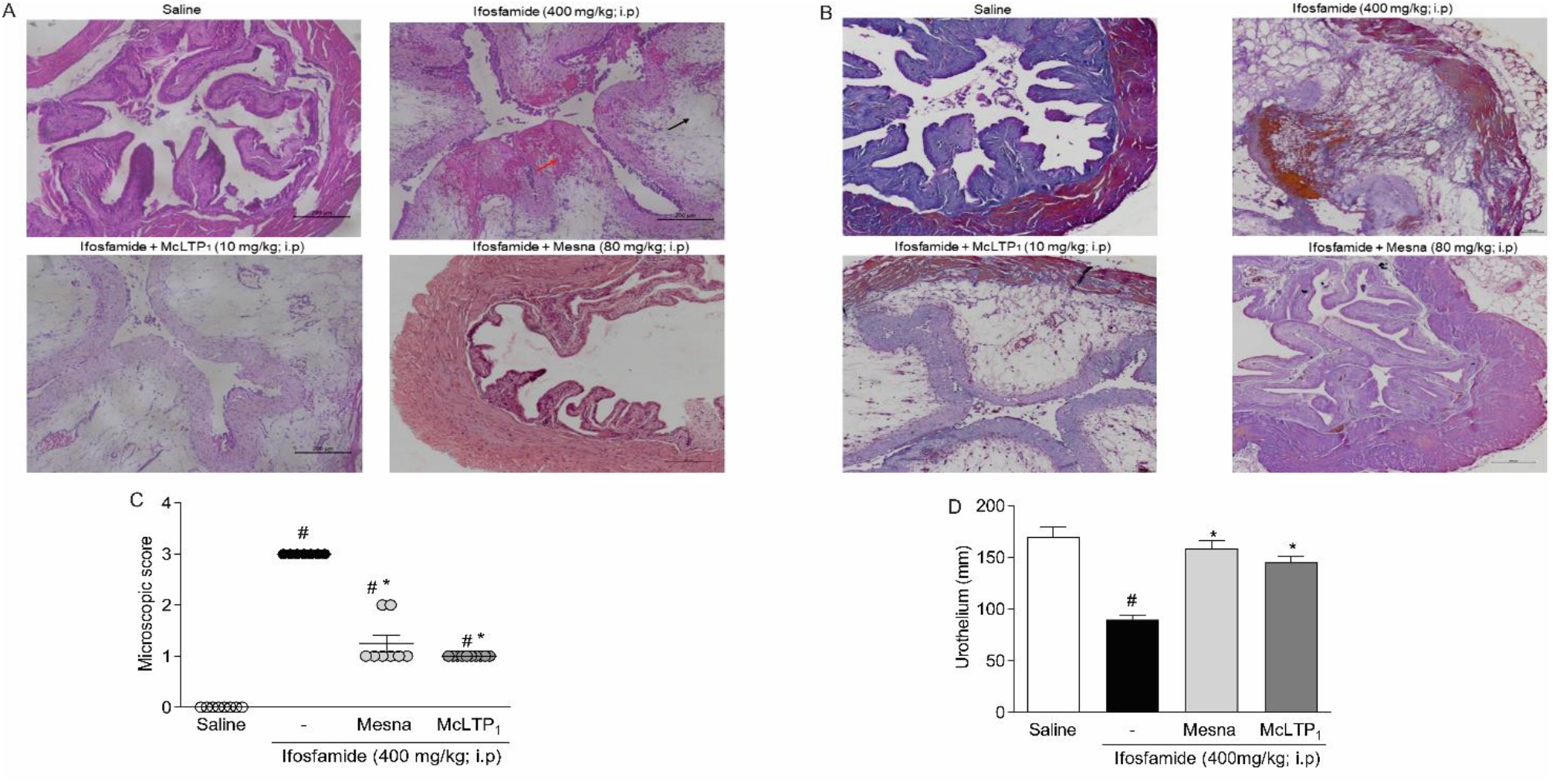
Effects of McLTP1 on histopathological changes in bladder tissues of mice with ifosfamide- induced hemorrhagic cystitis. Representative histological alterations of the bladders obtained from rats of different groups (hematoxylin and eosin staining, × 40 magnification). The bladders of IFO-injected mice displayed extensive area of edema (black arrow) and hemorrhage (red arrow). The results are reported as means ± EPM (n = 6-8/group). #P < 0.05 vs. saline group (negative control); *P < 0.05 vs. IFO group.

### Effect of McLTP_1_ on histopathological changes in ifosfamide-induced hemorrhagic cystitis

IFO induced significant histopathological injury (3[3-3]) compared to the saline group (0[0–0]) (Fig. 2C). Such damage was prevented in animals treated with McLTP_1_ (10 mg/kg) (1[1-1]) compared to IFO. These parameters were also attenuated by mesna, which was used as a positive control (1[1-2]). The injection of IFO induced a pronounced loss of epithelial cell lining, accentuated edema, hemorrhage, vascular congestion and ulceration in contrast to normal saline controls (Fig. 2A). The preventive effect of McLTP_1_ is also depicted in Fig. 2A, which shows mild edema, a preserved epithelial surface. The presence of hemorrhage was investigated with the special stain, where an intense staining is observed in hemorrhagic foci of bladders from the ifosfamide group, compared to a marked reduction after treatment with McLTP_1_ and mesna (Fig. 2B). Treatment with ifosfamide also induces a reduction in the thickness of the urothelium (89.73 ± 4.44 mm), when compared to the control group (169.4 ± 10.17 mm). McLTP_1_ and Mesna were able to preserve this epithelium that covers a large part of the urinary tract (145.6 ± 4.96 mm; 158.3 ± 7.84 mm), respectively (Fig. 2D).

### McLTP_1_ attenuated inflammatory parameters in ifosfamide-induced hemorrhagic cystitis

As shown in Fig. 3, IFO significantly increased TNF-α, IL-1ß and IL-6 tissue levels (TNF-α: 86.13 ± 9.51; IL-1β: 95.48 ± 4.77; IL-6: 32.37 ± 3. 03 pg/mg of tissue, respectively), but did not alter the levels of the anti-inflammatory cytokine IL-10 (6.99 ± 0.81 pg/mg of tissue), compared to saline injected animals (TNF-α: 13.0 ± 1 .07; IL-1β: 20.96 ± 1.41; IL-6: 12.39 ± 2.37; IL-10: 4.94 ± 1.22 pg/mg of tissue, respectively). In addition, McLTP1 reduced TNF-α, IL-1ß and IL-6 levels (TNF-α: 16.25 ± 1.32; IL-1β: 73, 20 ± 2.86; IL-6: 20.23 ± 1.96 pg/mg tissue) versus the IFO group. As well as, it promoted an increase in the release of the anti-inflammatory cytokine IL-10 (28.42 ± 5.47 pg/mg of tissue), similar to the treatment with mesna, except in the anti-inflammatory cytokine IL-10, where mesna did not increase significantly compared to the ifosfamide group (TNF-α: 19.21 ± 2.24; IL-1β: 63.20 ± 3.84; IL-6: 10.69 ± 2.13; IL-10: 14.92 ± 2.67 pg/mg tissue).

**Fig. 3.**
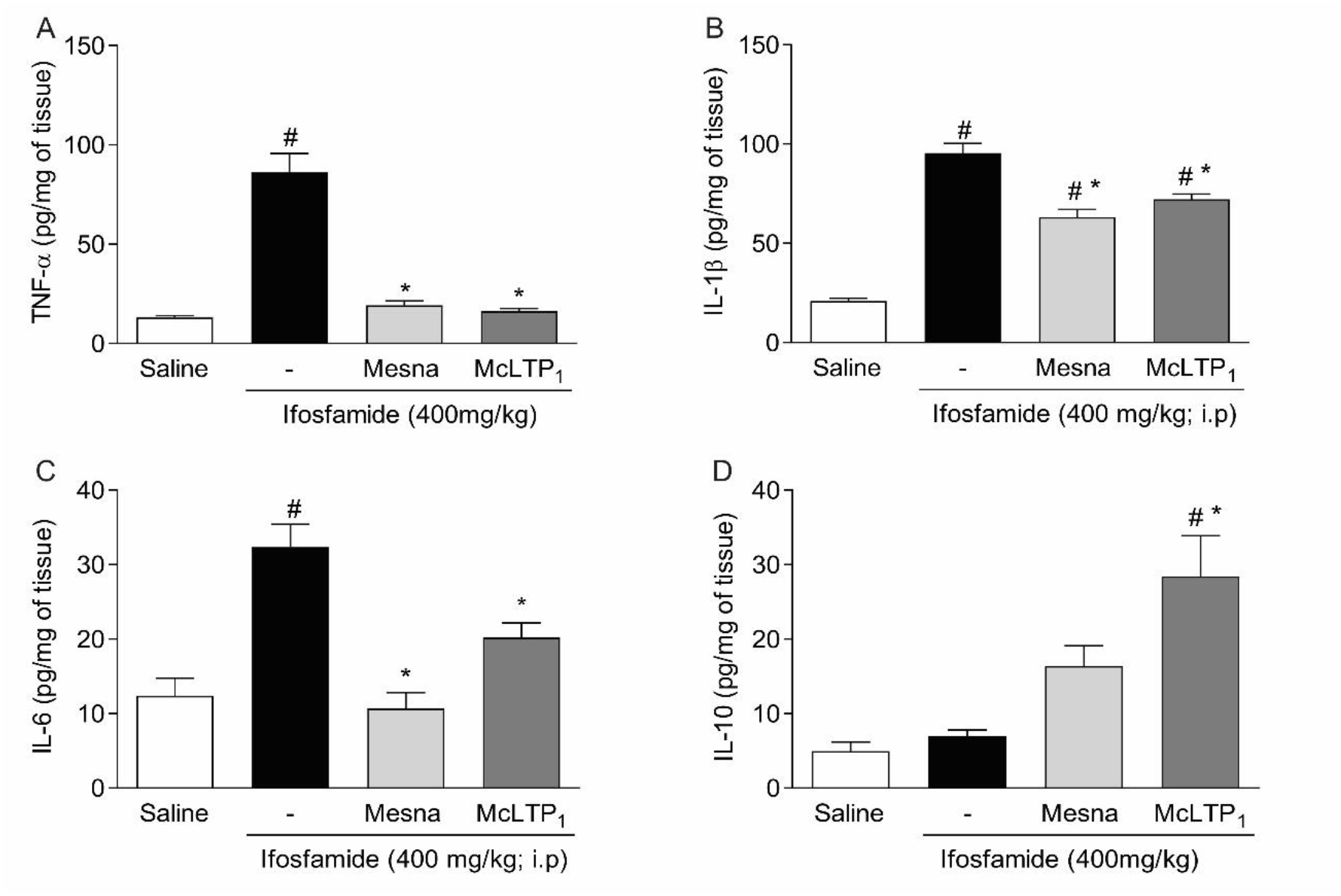
Effect of McLTP_1_ treatment on tissue levels of TNF-α, IL-1β, IL-6 and IL-10 in ifosfamide-induced hemorrhagic cystitis. The results (picogram/mg of tissue) were expressed as mean ± EPM. (n=8). # p<0.05 in relation to the saline group and * p<0.05 in relation to the group that received only ifosfamide.

**Fig. 4.**
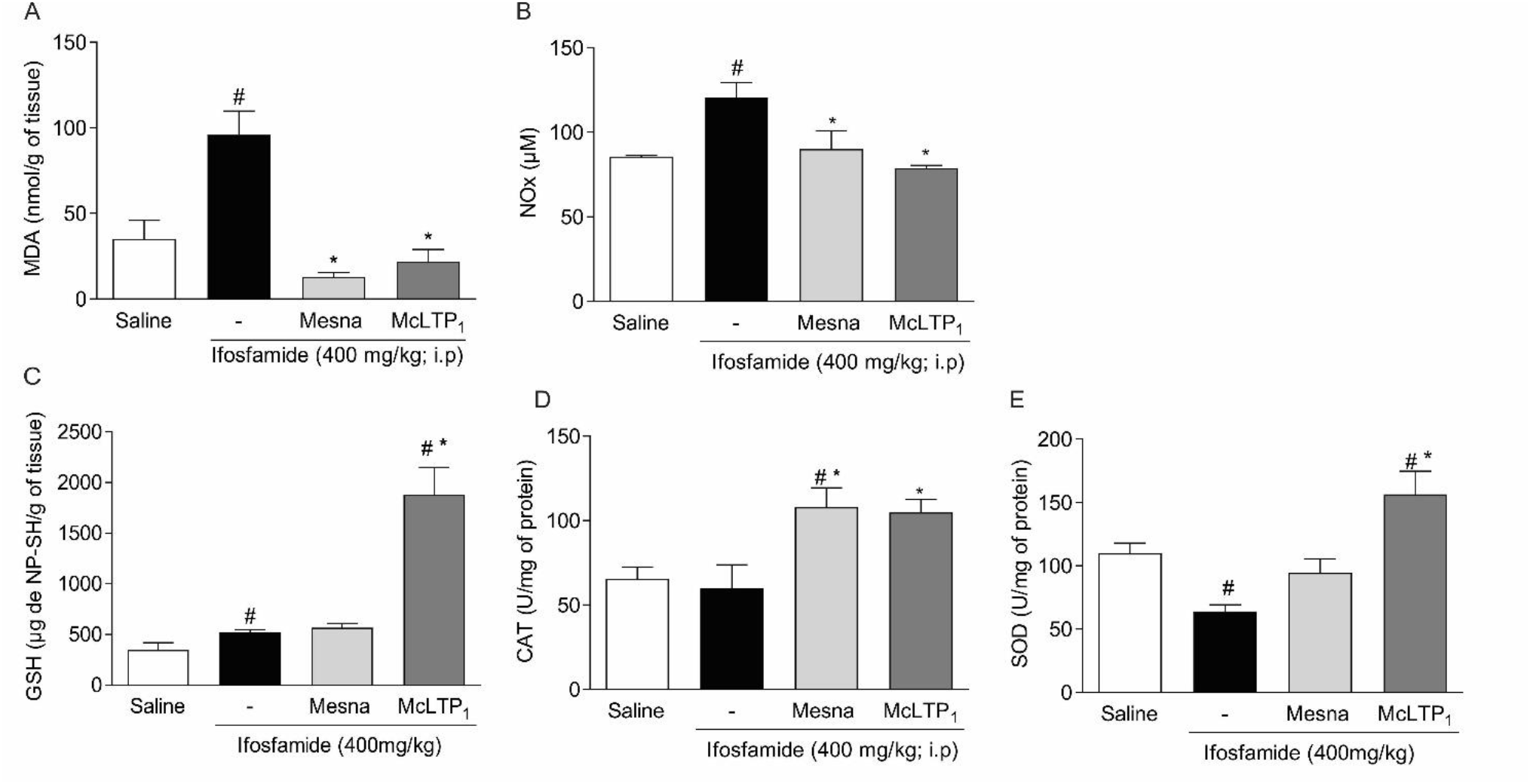
Effect of McLTP_1_ treatment on oxidative stress in ifosfamide-induced hemorrhagic cystitis. The oxidative stress markers MDA, Nitrite, SOD, CAT and antioxidant, GSH were measured in the bladder. Results were expressed as mean ± SEM. (n=8). # p<0.05 in relation to the saline group and * p<0.05 in relation to the group that received only ifosfamide.

### Effect of McLTP_1_ on oxidative stress in ifosfamide-induced hemorrhagic cystitis

The evaluation of oxidative stress was carried out through the indirect determination of Nitrite (NO), the degree of lipid peroxidation, through the marker malonaldehyde (MDA) and the antioxidant enzymes reduced glutathione (GSH), catalase (CAT) and superoxide dismutase (SOD). The saline group had MDA, NO, GSH, SOD and CAT levels determined in the bladder (MDA: 35.11 ± 10.78 nmol/g of tissue; NO: 85.57 ± 0.918 µM; GSH: 347 ± 73. 78 μg/g tissue; CAT: 65.59 ± 6.92 U/mg protein; SOD: 109.7 ± 8.23 U/mg protein). Oxidative stress increased significantly in the ifosfamide group, with high levels of lipid peroxidation determined by quantification of MDA (95.96 ± 13.90 nmol/g of tissue) and NO (120.5 ± 9.02 µM) compared to the group control. Also leading to depletion of antioxidant enzyme levels (GSH: 522.2 ± 27.23 μg/g of tissue; CAT: 59.85 ± 13.93 U/mg of protein; SOD: 63.94 ± 5.33 U /mg of protein). However, treatment with McLTP1 was effective in significantly increasing the activity of the antioxidant enzymes GSH, CAT and SOD in the tissue (GSH: 1906 ± 161.3 μg/g of tissue; CAT: 104.8 ± 7.79 U/mg of protein; SOD: 156.4 ± 11.09 U/mg of protein), as well as restoration of activity to normal values of NO (78.74 ± 1.74 µM) and lipid peroxidation (MDA: 21.79 ± 7.12 nmol/g tissue), similar to mesna treatment. However, the antioxidant activity of mesna was only evidenced in the catalase dosage, while there was no significant increase for GSH and SOD (GSH: 568.4 ± 41.80 μg/g of tissue; CAT: 108 ± 11.45 U/mg of protein; SOD: 94.48 ± 11.09 U/mg of protein; NO: 90.06 ± 10.84 µM; MDA: 12.84 ± 2.54 nmol/g of tissue) (Fig. 16).

### Effect of McLTP_1_ on COX-2 and TNF-α immunostaining in ifosfamide-induced hemorrhagic cystitis

IFO significantly increased COX-2 immunostaining (86.82 ± 7.24) and the relative expression of COX-2 mRNA (408.9 ± 91.84) (Fig. 5) compared to the saline group (Immunostaining: 3.22 ± 0.35; mRNA expression: 1.01 ± 0.36). Consistently, with its anti-inflammatory effect, McLTP1 reduced COX-2 (Immunostaining: 13.22 ± 1.65; mRNA expression: 31.33 ± 17.86) versus IFO-injected mice. Similar to that with mesna (Immunostaining: 14.90 ± 2. 59; mRNA expression: 6.39 ± 2.99). Figure 6 shows intense TNF-α staining in the ifosfamide group (67.34 ± 3.45) when compared to the saline negative control (4.52 ± 0.49), while treatment with McLTP_1_ significantly reduces immunostaining (22.92 ± 3.16), similar to treatment with mesna (26.69 ± 2.11), both significantly different from the group treated with ifosfamide alone.

**Fig. 5.**
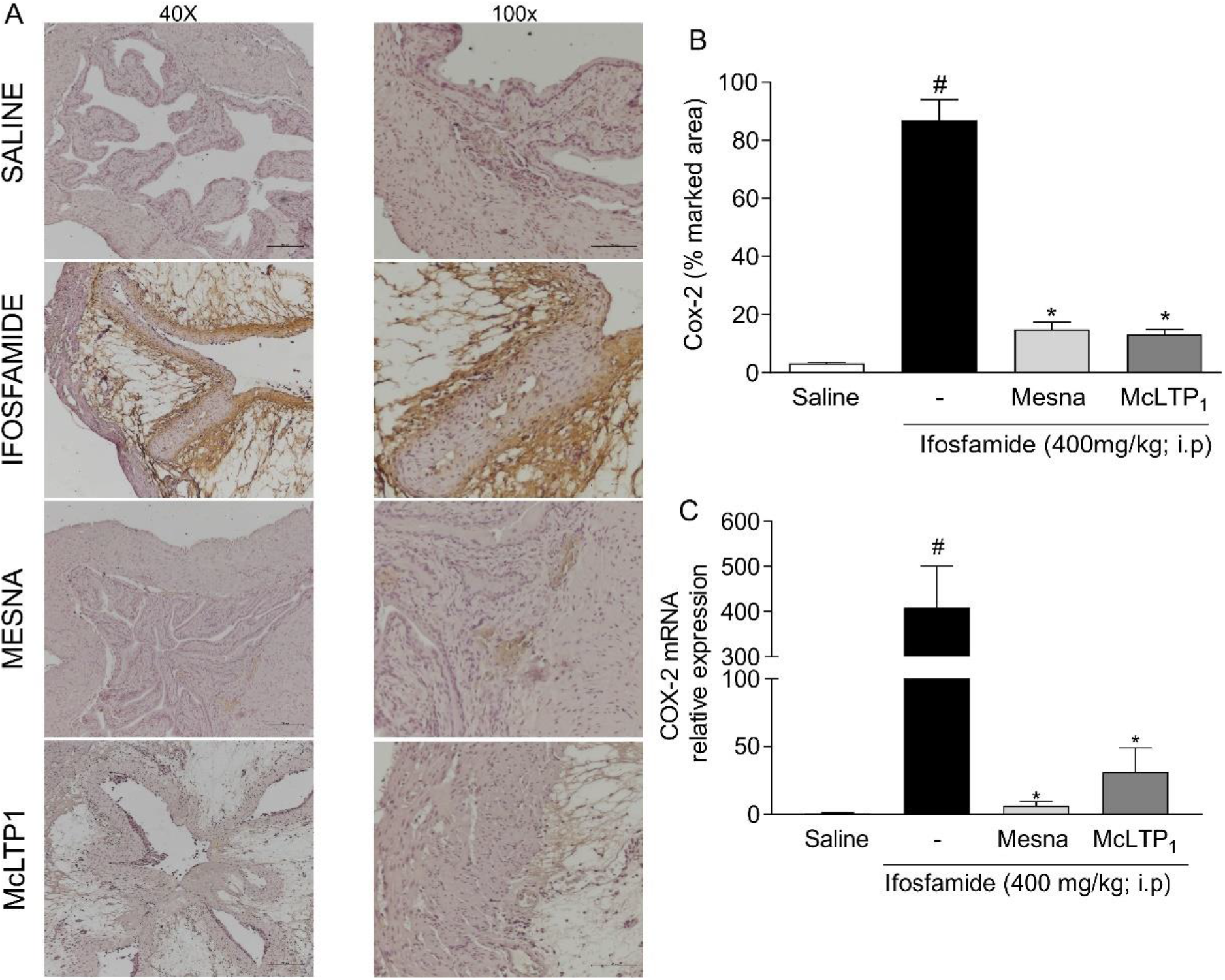
Effect of McLTP_1_ treatment on COX-2 immunostaining in ifosfamide-induced hemorrhagic cystitis. Results were expressed as mean ± SEM. (n=8). # p<0.05 in relation to the saline group and * p<0.05 in relation to the group that received only ifosfamide.

**Fig. 6.**
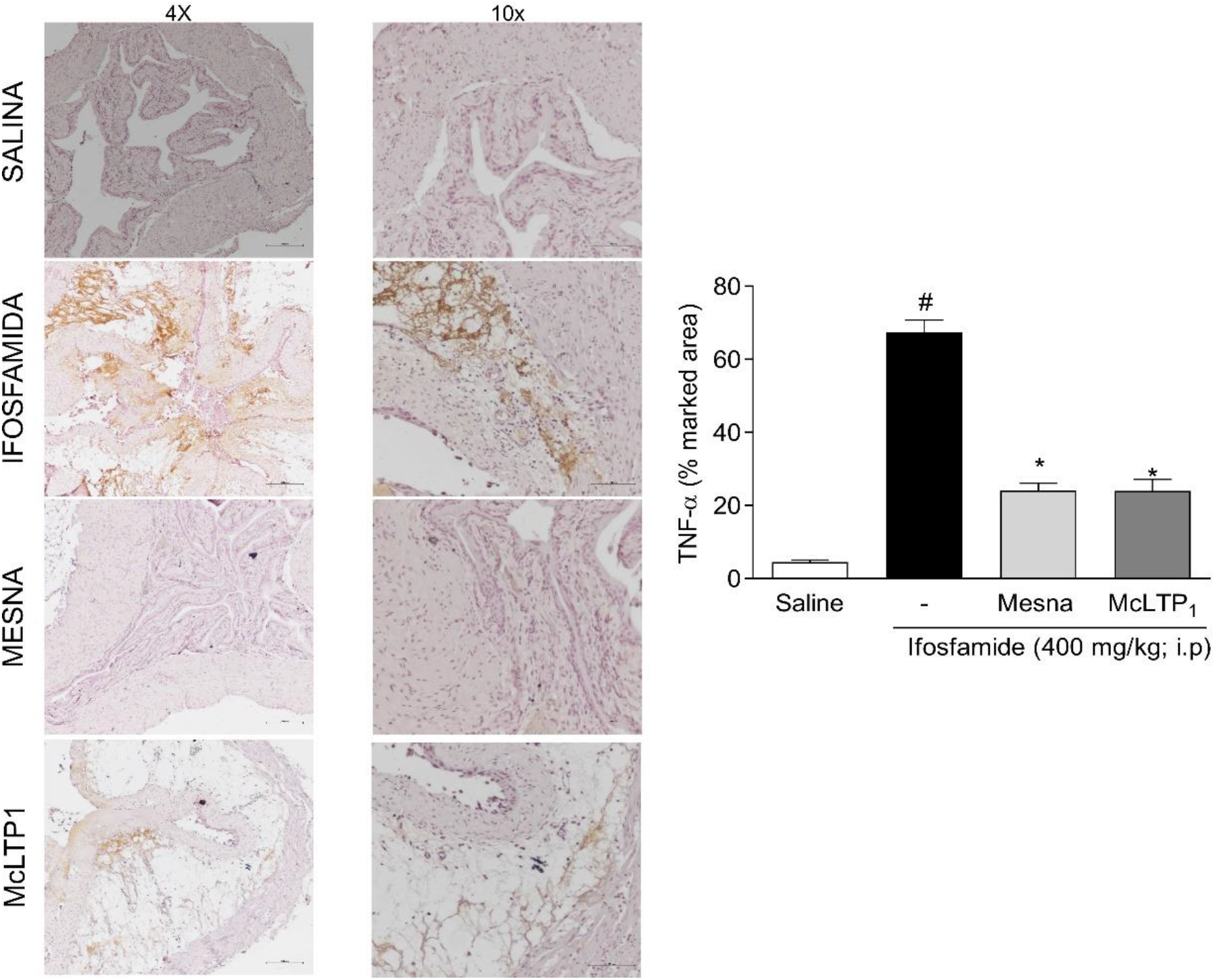
Effect of McLTP_1_ treatment on TNF-α immunostaining in ifosfamide-induced hemorrhagic cystitis. Results were expressed as mean ± SEM. (n=8). # p<0.05 in relation to the saline group and * p<0.05 in relation to the group that received only ifosfamide.

### Effect of McLTP_1_ on F4/80 and NF-kB immunofluorescence in ifosfamide-induced hemorrhagic cystitis

Ifosfamide significantly increased F4/80 expression (43.95 ± 6.53%) in the bladder when compared to the control group (2.13 ± 0.33%) (Figure 7). McLTP_1_ and Mesna were able to reduce F4/80 marking when compared to the ifosfamide group (5.46 ± 0.55%; 4.17 ± 0.55%), respectively. In immunofluorescence for NF-kB, ifosfamide significantly increased expression of this transcription factor (12.14 ± 1.28%), compared to the group that received only saline solution (control group) (1.53 ± 0.13%). McLTP_1_ and Mesna treatment were able to reduce NF-kB staining when compared to the ifosfamide group (3.22 ± 0.30 %; 2.18 ± 0.17 %) (Figure 8).

**Fig. 7.**
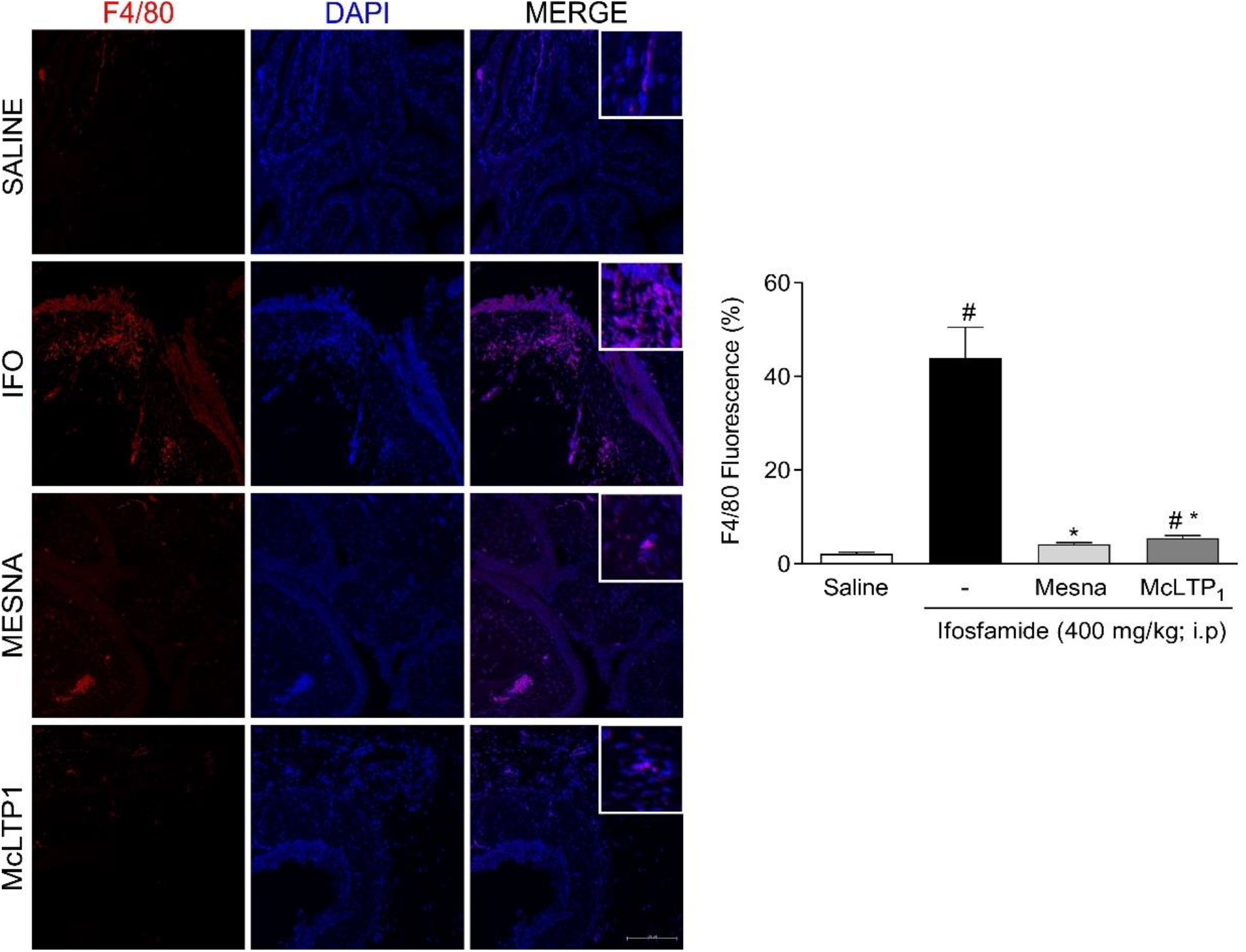
Effect of McLTP_1_ treatment on F4/80 immunostaining in ifosfamide-induced hemorrhagic cystitis. Values are presented as mean ± standard error of mean (SEM) of the percentage of positive fluorescent area of F4/80 expression compared to saline expression. *P < 0.05 versus control group. Source: survey data. Red: F4/80; blue: DAPI (nuclear marker). Increase: 200x. Scale: 50 µm.

**Fig. 8.**
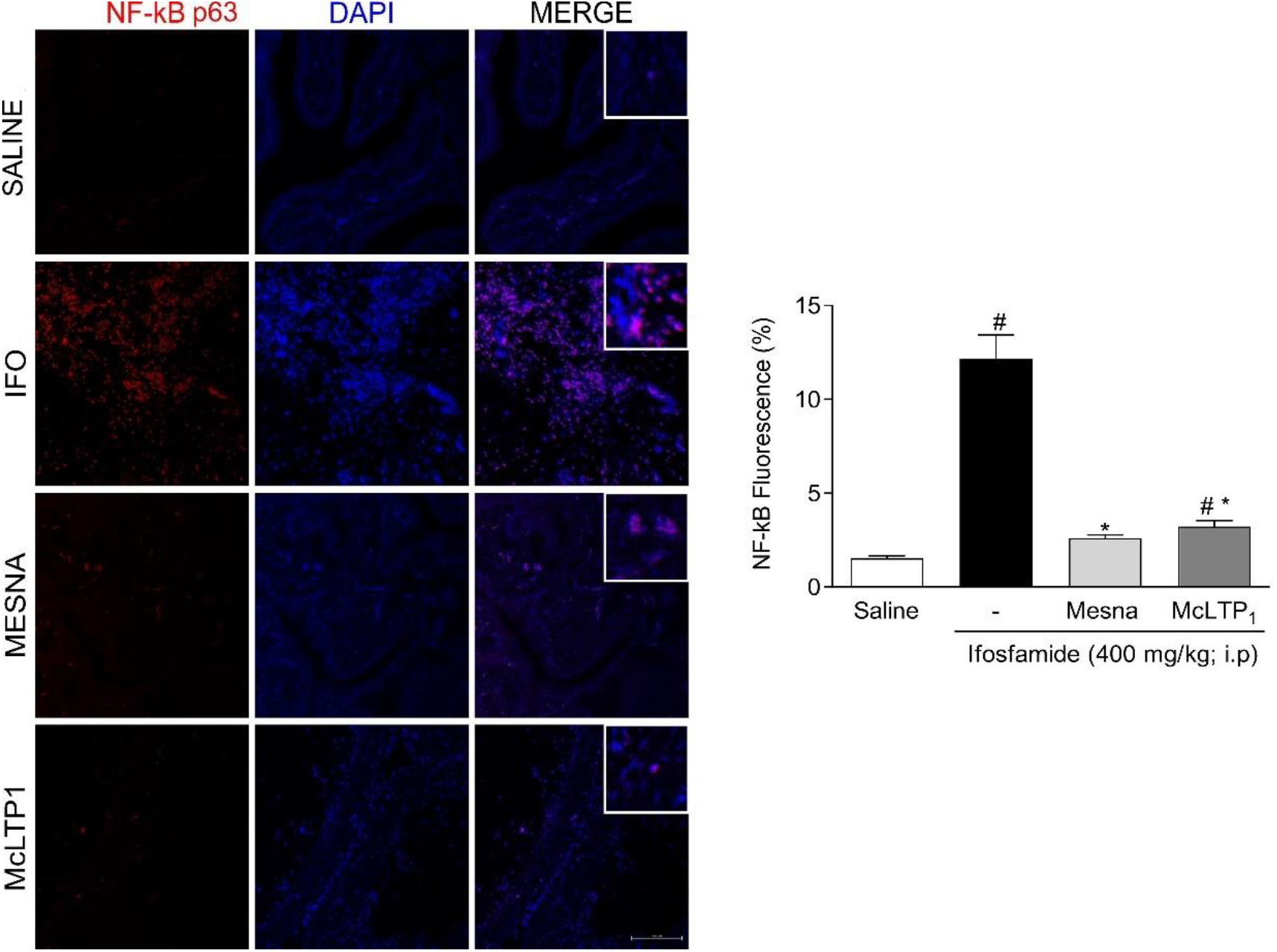
Effect of McLTP_1_ treatment on NF-kB immunostaining in ifosfamide-induced hemorrhagic cystitis. Values are presented as mean ± standard error of mean (SEM) of the percentage of positive fluorescent area of NF-kB expression compared to saline expression. *P < 0.05 versus control group. Source: survey data. Red: NF-kB; blue: DAPI (nuclear marker). Increase: 200x. Scale: 50 µm.

### Effect of McLTP_1_ on IL-4, IL-33, and iNOS gene expression in ifosfamide-induced hemorrhagic cystitis

Samples of the animals’ bladders after administration of ifosfamide or saline were used to determine the gene expression of important markers of the inflammatory process. Treatment with ifosfamide increased gene expression of IL-4 (SAL: 0.98 ± 0.40 vs. IFO: 4.18 ± 0.64), IL-33 (SAL: 0.36 ± 0.11 vs. IFO: 146.2 ± 47.46) and iNOS (SAL: 2.97 ± 1.12 vs. IFO: 24.65 ± 7.78) when compared to the saline group (P<0.05). However, treatment with McLTP_1_ reduced the expression of IL-4 (Mc: 0.67 ± 0.42 vs. Mesna: 0.40 ± 0.33), IL-33 (Mc: 5.90 ± 2.33 vs. Mesna: 1.18 ± 0.39), and iNOS (Mc: 1.29 ± 0.48 vs. Mesna: 0.67 ± 0.49), results similar to treatment with the standard drug mesna (Figure 9).

**Fig. 9.**
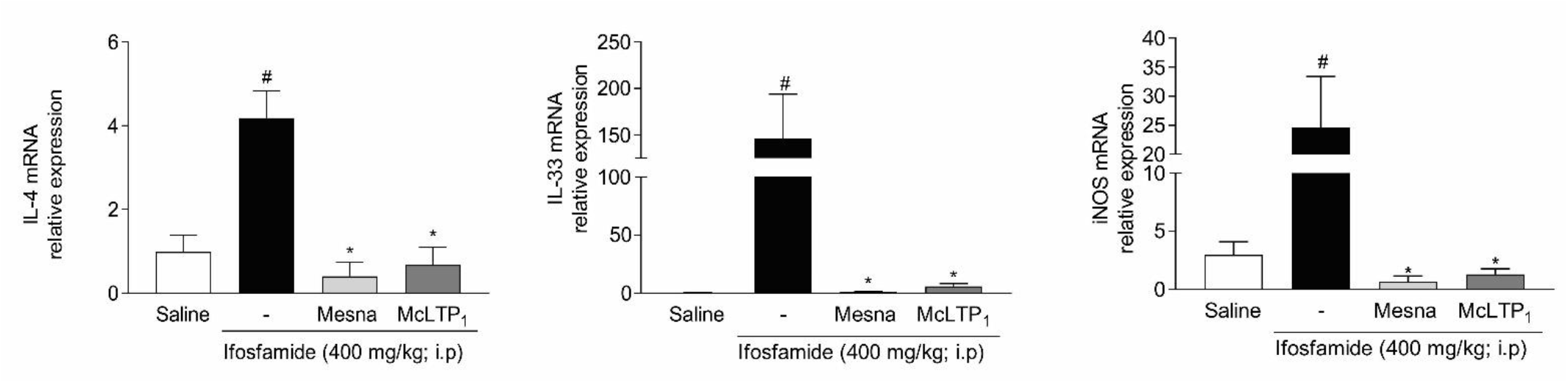
Effect of McLTP1 treatment on IL-4, iNOS and IL-33 gene expression in ifosfamide-induced hemorrhagic cystitis. Inflammatory markers were measured in the bladder. Results were expressed as mean ± SEM. (n=8). # p<0.05 in relation to the saline group and * p<0.05 in relation to the group that received only ifosfamide.

## Discussion

Chemotherapy with oxazaphosphorines, such as ifosfamide, is frequently limited by urotoxicity, in spite of its proven efficacy in malignant diseases, rheumatoid arthritis and systemic lupus erythematosis, IFO has several side effects and the main one that restricts clinical use is hemorragic cystitis [22]. The urotoxic mechanism this adverse effect involve the inflammatory process and oxidative stress, activating transcription factors of proinflammatory cytokine such as TNF-α and IL-1β, nitric oxide, migration of inflammatory cells such as macrophages and neutrophils, resulting histologically in subepithelial edema, ulceration, hemorrhage, and endothelial tissue destruction (Brock et al. 1979; Theman et al. 1987; Gonzalez et al. 2005; Macedo et al. 2012; Silva Junior et al. 2013), (Haldar et al. 2014). In this way, antioxidants and antiinflammatory agents may suppress its urotoxicity leading to improve the accession of treatment with IFO [1, 24].

In the present study, McLTP_1_ demonstrated anti-inflammatory and antioxidant activity in the model of IFO-induced cystitis, through attenuation of edema and hemorrhage, by reducing pro-inflammatory cytokines (TNF-α, Il-1β, IL-6, IL-33) and positively modulating the anti-inflammatory cytokine IL-10. As well as inhibiting the expression of important transcription factors such as NF-kB, COX-2 enzyme, iNOS and F4/80 macrophage. In addition, it reduced oxidative stress markers, MDA and nitrite, and was even able to increase antioxidant activity through upregulation of enzymes, SOD, catalase and GSH.

Previous studies with McLTP_1_ described in the literature already show its anti-inflammatory and antioxidant activity in different preclinical models of nociception, intestinal mucositis by irinotecan, antimicrobial and protective activity on sepsis, nephro and gastroprotective. Therefore, the protein was considered a potential uroprotector to be investigated [42–45].

A pilot study with the dose-response curve was carried out by administering McLTP1 at doses of 10, 20 or 40 mg/kg intraperitoneally to evaluate macroscopic and basic inflammatory parameters. A group treated with mesna (80 mg/kg; i.p.) was also applied because it is the drug currently used by patients undergoing treatment with oxazaphosphorines [13].

The formation of edema which is characteristic of HC was observed indirectly through the evaluation of the mean bladder wet weight, that both Mesna and McLTP_1_ showed that they were capable of preventing the appearance of edema. However, macroscopic scores showed that only the lowest dose (10mg/kg) was able to reduce edema and hemorrhage. From the onset of inflammation in the bladder, the activity of the myeloperoxidase enzyme was measured, as an indirect way to determine the neutrophilic infiltration [10]. Ifosfamide increased tissue MPO levels, which were significantly reduced after treatment with McLTP_1_ at doses of 10 and 20 mg/kg. Our findings are in line with previous studies, in which McLTP_1_ (8mg/kg) reduced MPO activity in the duodenum of mice subjected to intestinal mucositis by irinotecan [42]. In fact, previous research with other models of inflammation has already shown that polysaccharides from Morinda citrifolia are able to attenuate classic signs of inflammation such as edema in a model of experimental colitis induced by acetic acid [46]. That said, the lowest protein dose (10 mg/kg) was chosen to continue the study.

In order to quantify more precisely the extent of edema and hemorrhage, permeability by Evans Blue and measuring hemoglobin were investigated in bladder tissue [47]. McLTP_1_ reduced vascular permeability and consequently release of protein-bound fluids (albumin) from blood to tissues, as well as bladder hemoglobin levels when compared to ifosfamide treatment. However, in the mesna group there was a reduction only in hemoglobin and it was not able to prevent the formation of edema by the aforementioned method.

McLTP_1_ reduced the severity of histological changes, such as edema, hemorrhagic foci, vascular congestion, inflammatory infiltrate and urothelial denudation. The protein stood out for preserving the urothelium and controlling hemorrhage. Therefore, the measurement of the thickness of the urothelium was carried out in order to confirm this finding, where McLTP_1_ and mesna presented measurements similar to the negative control. Moreover, hemorrhagic cystitis induced by oxazaphosphorine is characterized by massive extravasation of mature red blood cells and reticulocytes, marked by the presence of hematuria. This parameter was approached quantitatively by measuring tissue hemoglobin, which is directly proportional to vascular ruptures in bladder damage caused by ifosfamide [48]. The results obtained in this study show that McLTP_1_ had a protective effect against bladder hemorrhage by preventing hemorrhage. In addition, the hemorrhagic foci can be evidenced in Masson’s staining, where they were presented in red, which was significantly reduced after treatment with the protein [49].

Concerning the inflammatory process, the urotoxic effect of acrolein leads to the progression of urothelial damage by stimulating inflammation in the connective tissue and epithelial cells, in addition to the migration of macrophages, carrying inflammatory cytokines to the site of injury. STAT3 and NF-κB pathways, both major activators of the inflammation and immune response cascade through the IL-1β/TNF-α /IL-6 triad, were upregulated after treatment with ifosfamide [11]. Furthermore, IL-6 also has the ability to regulate iNOS activation in the urinary bladder under acrolein-induced inflammatory condition [50]. In our study, McLTP_1_ was able to modulate these crucial cytokines of the inflammatory cascade. Corroborating with our data, a polysaccharide fraction of Morinda citrifolia L. reduced the release of pro-inflammatory cytokines IL-6, IL-1β, and TNF-α in macrophage cell culture, RAW 264.7 [51]. Specifically, McLTP_1_ was able to reduce TNF-α, IL-1β in the model of paw edema induced by carrageenan in mice [52].

Bladder injury can also activate adjacent mast cells through the release of the pro-inflammatory cytokine IL-33, which acts as an alarmin released by endothelial, epithelial and smooth muscle cells, and is therefore considered a molecular pattern associated with damage (DAMP). The mast cell and IL-33 activation axis was the central mechanism in LL-37-induced bladder inflammation and pain in the murine interstitial cystitis model [53]. While the involvement of this cytokine specifically in a model of hemorrhagic cystitis was demonstrated in an unprecedented way in our study, where ifosfamide increased its gene expression and McLTP_1_ was able to modulate the release of IL-33 [54,55].

A cytokine also capable of modulating the inflammatory process is IL-4, having demonstrated that 4 hours after induction of cystitis by cyclophosphamide there was a significant increase in mRNA, which persisted 48 hours later, similar to that found after treatment with ifosfamide [56]. Corroborating the findings of the present study, where ifosfamide increased IL-4 gene expression, being significantly reduced by McLTP_1_ [27].

The anti-inflammatory cytokine IL-10 controls inflammatory processes by suppressing the expression of pro-inflammatory cytokines. Clarke et al. (1998) believe that IL-10 attenuates TNF-α receptor expression and mediates anti-inflammatory effects through inhibition of the NF-kB transcription factor, a key secondary messenger for inducing the expression of genes of pro-inflammatory cytokines [57,58]. Associated with the reduction of pro-inflammatory cytokines, McLTP_1_ was able to increase the synthesis of IL-10, confirming the contribution of this protein in the modulation of the inflammatory process. Corroborating our study, McLTP_1_ increased IL-10 in the ethanol-induced gastric injury model [44].

The binding of IL-1β with its receptors induces the phosphorylation of kinases leading to the translocation of NF-κB to the nucleus leading to the expression of iNOS and COX-2 [59]. In the ifosfamide hemorrhagic cystitis model, it was shown that after inhibition with eterocoxib, a COX-2 inhibitor, there was a significant reduction in PGE2, reduced expression of COX-2 and macroscopic and microscopic parameters [5,6]. In the present study, treatment with McLTP_1_ reduced gene expression and immunostaining for COX-2. A study with ethanolic extract extracted from the fruit of Morinda citrifolia observed a decrease in COX-2 expression in cultured human gastric epithelial cells (AGS) during Helicobacter pylori infection [60].

One of the crucial mechanisms of inflammatory events after acrolein enters the uroepithelial cells is the effect on NF-κB [61], because when translocated to the nucleus, it optimizes the transcription of specific genes that regulate responses inflammatory cells, such as inflammatory cells and pro-inflammatory cytokines [62]. The involvement of this protein in hemorrhagic cystitis was demonstrated after induction of the model with cyclophosphamide in rats, participating as an important inflammatory mediator [63]. In that study, ifosfamide increased NF-κB expression and McLTP_1_ was able to significantly reduce its nuclear immunostaining. Corroborating these findings, Damnacanthal, an anthraquinone isolated from the root of Morinda citrifolia, inhibited NF-κB activation in mast cells stimulated with calcium ionophore [64].

Still in the context of the inflammatory process, nitric oxide (NO), a reactive free radical that is involved in inflammation, is involved in the pathogenesis of hemorrhagic cystitis induced by cyclophosphamide through the induced form of iNOS [65]. iNOS expression is also stimulated by NF-κB and COX-2 [66]. Therefore, we demonstrated that McLTP_1_ was able to reduce iNOS gene expression when compared to the group treated only with ifosfamide. Previous studies also showed that noni juice polyphenols inhibited iNOS and COX-2 expression in the liver of high-fat hamsters [67].

The F4/80 protein, a glycoprotein that has been widely used as a macrophage marker, since this monoclonal antibody was identified to detect antigens present exclusively in phagocytes [68]. Previous studies have shown the involvement of this important marker in the hemorrhagic cystitis model, where cyclophosphamide markedly increased macrophage infiltration in the urinary bladder [33]. In the present study, McLTP_1_ was able to reduce the labeling to F4/80 in the ifosfamide-induced hemorrhagic cystitis model.

In addition to inflammatory activity, oxidative stress participates in the pathogenesis of hemorrhagic cystitis. Lipid peroxidation was observed after administration of oxazaphosphorines through the increase of malondialdehyde in the bladder, as well as the downregulation of antioxidant agents (GSH, SOD and CAT), revealing oxidative damage. In this study, treatment with McLTP_1_ significantly reduced MDA compared to animals treated with ifosfamide alone. Similar effects were found with polysaccharides extracted from noni in mice on a high-fat diet [69]. The antioxidant defense occurs through the antioxidant enzymes glutathione, catalase and SOD, with a function mainly in combating the propagation of damage caused by reactive oxygen species [70]. In the present study, we observed a significant reduction of glutathione in animals with hemorrhagic cystitis induced by ifosfamide. While after treatment with McLTP_1_ there was an increase in these antioxidant agents. Corroborating this study, the protein showed antioxidant activity in the model of nephrotoxicity induced by gentamicin in rats [43].

According to the results presented above, McLTP_1_ obtained a protective effect against ifosfamide-induced hemorrhagic cystitis in Balb/c mice. Such activity was evidenced by the reduction of edema and hemorrhage (scores of macroscopic evaluation, bladder wet weight, vascular permeability, hemoglobin dosage), preservation of morphology (histopathological scores and measurement of urothelium), reduction of inflammation (decrease of myeloperoxidase, pro cytokines -inflammatory, enzymes and growth factors) and increased antioxidant activity (dosage of MDA, Nitrite, GSH, SOD and Catalase). This suggests that McLTP_1_ can be the target of genetic improvement through the creation of a recombinant protein, making its prospection in toxicity tests in patients and clinical trials feasible.

## Author contributions (mandatory) for all authors

(G.F.P. Rangel; H.D.Oliveira; N.M.N. Alencar; T.F.G. de Sousa) conceived or designed the study; (N.M.N. Alencar; H.D.Oliveira; R.C.P. Lima Júnior; G.F.P. Rangel; D.V.T. Wong; R.C.F. Leitão) analyzed the data; (G.F.P. Rangel; N.M.N. Alencar) wrote the paper; (G.F.P. Rangel; A.G.Cajado; L.M.A. Rabelo; A.F. Pereira; A.S. Costa) performed research.

